# BOS1 is a positive regulator of wounding induced cell death and plant susceptibility to *Botrytis*

**DOI:** 10.1101/2022.01.18.476848

**Authors:** Fuqiang Cui, Xiaoxiao Li, Wenwu Wu, Wenbo Luo, Ying Wu, Mikael Brosché, Kirk Overmyer

## Abstract

Programmed cell death (PCD) is required for many aspects of plant biology, including stress responses, immunity, and plant development including root and flower development. Our understanding of PCD regulation is incomplete, especially concerning regulators involved in multiple divergent processes. The *botrytis-suscetible1* (*bos1*) mutant is one of the genotypes most susceptible to *Botrytis cinerea* (*Botrytis*) and has revealed the role of *BOS1* in cell death propagation during plant responses to wounding. The *bos1-1* allele harbours a T-DNA located in the 5’UTR upstream from the start codon that results in elevated *BOS1* transcript levels. Here, we resequenced the *bos1-1* genome and found a *MAS* promoter at the ends of the T-DNAs. Expression of the *BOS1* gene under control of the *MAS* promoter conferred the characteristic *bos1-1 Botrytis-* sensitivity and wounding phenotypes in wildtype plants. We used Crispr-Cas9 to create new *bos1* alleles that disrupt exons. These lines lacked the typical *bos1-1* wounding *and Botrytis* phenotypes, but exhibited reduced fertility, as previously observed in other *bos1* T-DNA alleles. With multiple overexpression lines of *BOS1*, we demonstrate that *BOS1* is involved in regulation of cell death propagation in a dosage dependent manner. Our data support that *bos1-1* is a gain-of-function mutant and that BOS1 acts as a positive regulator of wounding and *Botrytis*-induced PCD. Taken together these finding suggest that BOS1 function in both fertility and *Botrytis* response could be unified under cell death control.

## Introduction

Programmed cell death (PCD) is a finely tuned process, which occurs for example during plant-pathogen interactions and plant development, and has three stages including cell death initiation, propagation, and containment (McCabe, 2013; Van Hautegem et al., 2015). Many PCD regulators have been identified in *Arabidopsis thaliana* (*Arabidopsis*) from lesion mimic mutants that spontaneously develop cell death (Lorrain et al., 2003; Bruggeman et al., 2015). A separate class of regulators have been recognized from so called propagation class lesion mimic mutants, in which uncontained or “runaway” spreading cell death that can consume the entire leaf is observed once cell death is initiated (Lorrain et al., 2003). Although these have been crucial to understand the processes involved in regulation of cell death, for example the central role of several plant hormones (Bruggeman et al., 2015), our understanding of the signals leading to propagation and containment of cell death remains incomplete. Uncontained abscisic-acid dependent PCD propagation was found in *botrytis-suscetible1-1* (*bos1-1*; Cui et al., 2013), a mutant allele of *BOS1/MYB108* (At3g06490; Mengiste et al. 2003). PCD propagation was enhanced in *bos1-1* once death was initiated by pathogen infection or simply mechanical injury (Cui et al., 2013, 2019). Mechanical injury results in local cell death immediately adjacent to the wound in order to re-establish the integument (McCabe, 2013; Cui et al., 2013; Bostock and Stermer, 1989; Iakimova and Woltering, 2018). This wound-induced cell death response allows controlled PCD development at a fixed site, which makes wounding of *bos1-1* an attractive experimental system for studies on PCD propagation (McCabe, 2013). BOS1 is a R2R3 MYB transcription factor, which was first functionally characterized with the *bos1-1* mutant, based on its striking susceptibility to the necrotrophic fungal pathogen *Botrytis cinerea*, in the seminal paper by Mengiste et al. (2003). Subsequently, *BOS1* has been recognized as a key gene involved in plant-pathogen interactions, and *bos1-1* has helped to reveal the important role of cell death in susceptibility to necrotrophic fungi (Kraepiel et al., 2011; Cui et al., 2013, 2019). The *bos1-1* allele was genetically characterized as a recessive loss-of-function mutant, although the T-DNA insertion is located in the 5’UTR upstream from the start codon and results in abnormally high expression of *BOS1* (Mengiste et al., 2003).

PCD is also indispensable for plant development, including reproductive development where PCD is required for both proper development and release of pollen (Mandaokar and Browse, 2009; Daneva et al., 2016; Xu et al., 2019). One open question in plant PCD research is to which extent pathogen activated and developmental PCD overlap in regulatory mechanisms and execution (Huysmans et al., 2017). BOS1 functions in both plant stress responses and development (Mandaokar and Browse, 2009; Kraepiel et al., 2011; Cui et al., 2013; Xu et al., 2019; Cui et al., 2019). BOS1 can be ubiqutinated by BOTRYTIS SUSCEPTBLE1 INTERACTOR (BOI), an E3 ligase that attenuates stress induced cell death in plants (Luo et al., 2010). Three mutant alleles with T-DNA insertions in the first intron were used to study the role of *BOS1/MYB108* in anther development (Mandaokar and Browse, 2009). The mutants displayed reduced male fertility, lower pollen viability, and delayed anther dehiscence. However, the stress response of these alleles remains untested. As different *bos1* alleles were used in the study of pathogen versus developmental PCD, there is a lack of information to which extent this transcription factor could act in both types of cell death. Further, the existing mutants for *bos1* are either intron insertions (Mandaokar and Browse, 2009) or a 5’-UTR insertion (*bos1-1*; Mengiste et al., 2003), which makes interpretation of BOS1 function in cell death control unclear. Here we generated new *bos1* exon mutant alleles and present evidence that this transcription factor is a positive regulator of cell death, in contrast to the roles previously assigned to BOS1.

## Results

### *Botrytis* and wounding response in new *bos1* Crispr alleles

Genome editing allows the generation of desired mutants (Jiang et al., 2013; Xing et al., 2014). We used CRISPR-Cas9 to create three *BOS1* loss-of-function alleles, targeting the first and second exons (*bos1-c1* to -*c3*; Fig. 1A). These mutations caused frame-shifts resulting in truncated proteins (Fig. 1B). None of these alleles exhibited the characteristic *bos1-1* phenotypes seen with *Botrytis* infection or wounding treatments. *Botrytis*-induced lesion sizes and wound-induced cell death spread in these mutants were similar to wildtype (Fig. 1C and D). This suggested that *bos1-1* may be not a true loss-of-function mutant of *BOS1*. To confirm this hypothesis, we generated the heterozygous mutant *bos1-1/BOS1* by a cross between *bos1-1* and wildtype. Upon wounding and *Botrytis* treatments, *bos1-1/BOS1* exhibited intermediate phenotypes between wildtype and the *bos1-1* homozygous mutant; both the extent of wounding-induced runaway cell death and the size of *Botrytis*-induced lesions in *bos1-1/BOS1* were significantly larger than wildtype but smaller than in *bos1-1* (Fig. 1C and D). Further, we tested the distribution of *Botrytis-* and wounding-induced phenotypes in a F_2_ population derived from a confirmed *bos1-1/BOS1* heterozygote F_1_ individual. These phenotyped F_2_ populations revealed a 1:2:1 segregation ratio, fitting the model where *bos1-1* is a co-dominant gain-of-function allele; in which one quarter exhibited *bos1-1* symptoms; one half had intermediate phenotypes, similar to *bos1-1/BOS1;* and one quarter had wild type characteristics (Supplemental Fig. S1). Plants of one replicate were genotyped, which confirmed that the plants exhibiting enhanced *Botrytis-induced* lesion size were *bos1-1* homozygotes while the population exhibiting intermediate sized lesions were *bos1-1/BOS1* heterozygotes (Supplemental Fig. S1). The genetics of both the F_1_ and F_2_ generations supports that *bos1-1* is a co-dominant gain-of-function mutant.

**Figure 1.**
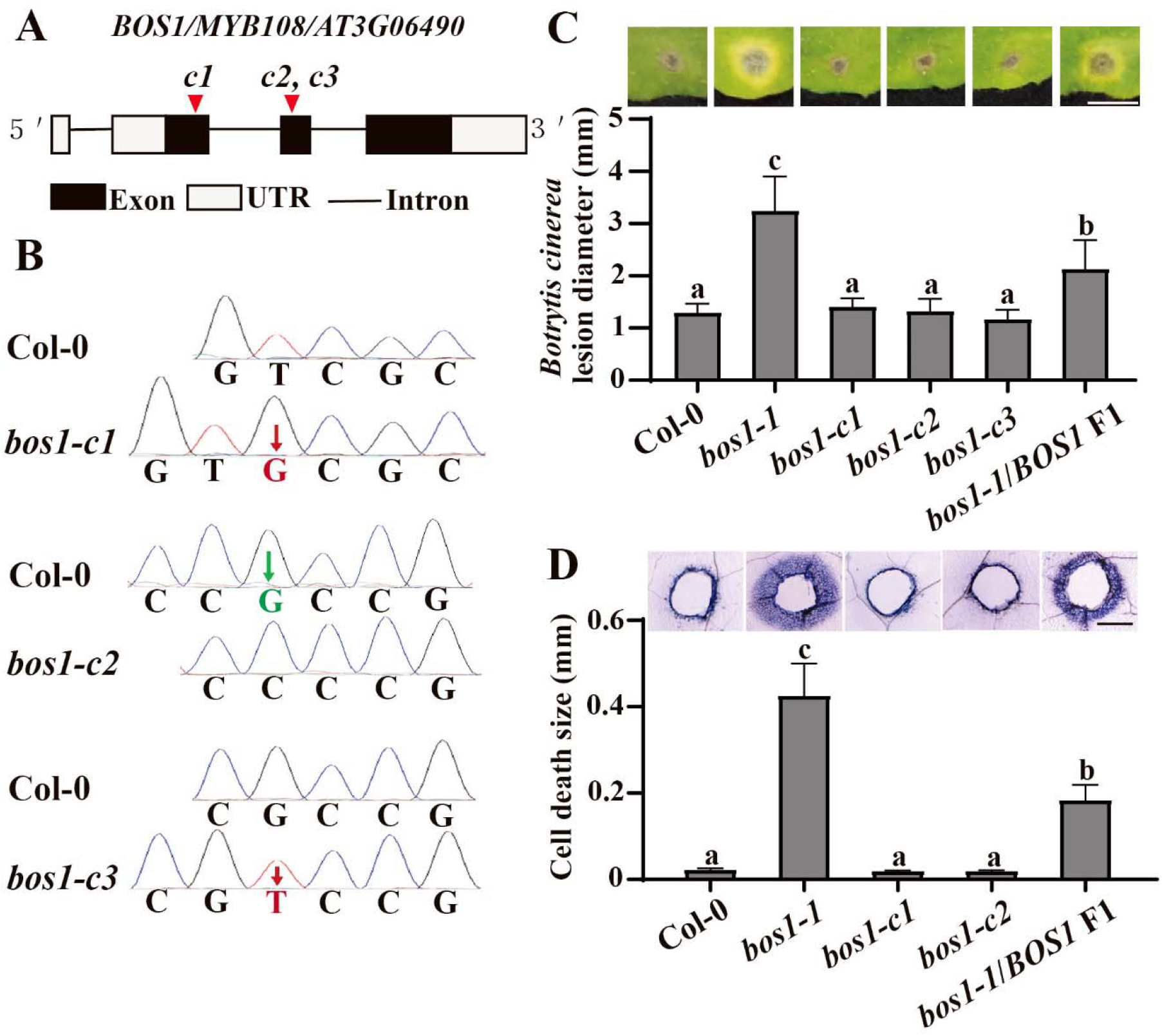
New *bos1* alleles created with Crispr-Cas9 did not exhibit *bos1-1* phenotypes. **(A)** Schematic diagram of the new *bos1* insertion and deletion alleles. The guide RNA (gRNA) positions are indicated with red triangles and the new *bos1* alleles made by CRISPR were designated as *c1, c2* and *c3*. **(B)** Genome editing induced changes in the indicated mutants. Single base insertions (red characters) and deletion (green characters) were detected with Sanger sequencing. **(C)** Disease symptoms and lesion sizes induced by inoculation with *Botrytis cinerea*. Droplets of conidia suspension (3 μl, 2×10 spores ml^-1^) were inoculated on fully expanded leaves of the indicated genotypes. Symptoms were photographed at three days post inoculation. Lesion sizes were measured in ImageJ and statistically analysed with one-way ANOVA (three independent biological replicates; *n*=24 in total). Scale bar=0.5 cm. **(D)** Wound induced cell-death spread. Leaves punctured with a toothpick were subjected to trypan blue staining to visualize dead tissue at four days post wounding. Representative pictures are shown to illustrate the dead tissues around the wounds. The extent of cell death spread was measured from the edge of the wound to the outer frontier of spreading cell death. These experiments were performed three times with similar results (*n*=24 in total). Scale bar=0.5 mm.

### Crispr *bos1* alleles are impaired in fertility

The CRISPR knock-out lines exhibited strong deficiencies in fertility, which was observed as the number of siliques with reduced size and a delay in flower senescence (Fig. 2A). This finding is consistent with the reduced fertility phenotypes of mutants with T-DNA in the introns of *BOS1* (Xu et al., 2019; Mandaokar and Browse, 2009). Previous studies suggested that the impaired fertility of *bos1* intron T-DNA alleles was due to deficient or delayed pollen release, as their anthers were mostly still closed (Xu et al., 2019). We examined the anthers of the new CRISPR knock-out alleles and found the same phenotype. Pollen release was significantly reduced or delayed in comparison to wild type (Fig. 2B and C). In contrast, *bos1-1* did not exhibit any phenotypes in fertility and anther dehiscence (Fig. 2). The phenotypic similarity between our CRISPR knock-out lines and the previously used *BOS1* intron T-DNA alleles further support that *bos1-1* is not a loss-of-function mutant of *BOS1*.

**Figure 2.**
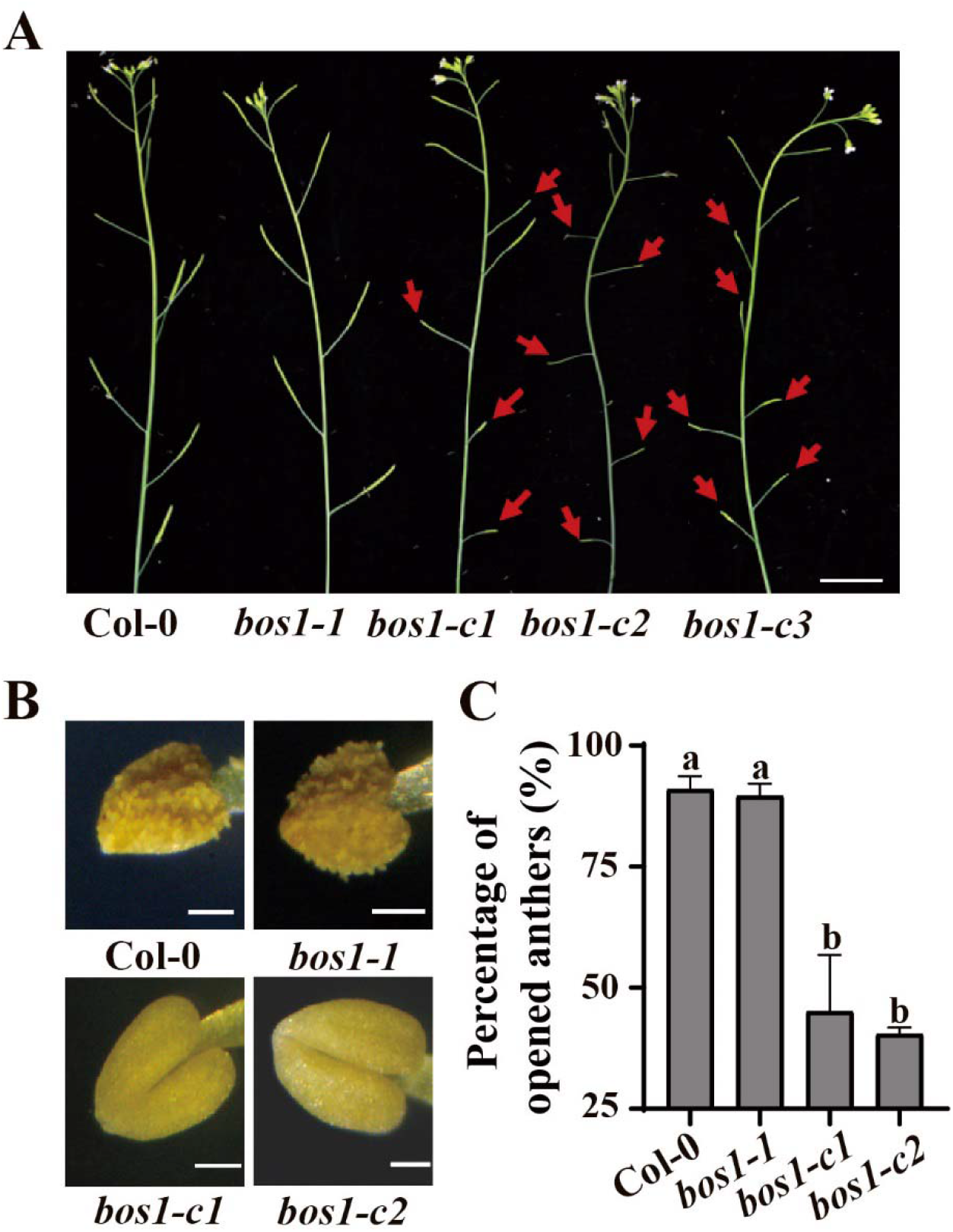
Crispr *bos1* knock-out lines were impaired in pollen release. **(A)** The *bos1* alleles created with Crispr-Cas9 resulted in impaired fertility. Red arrows indicate siliques with reduced seed production. Delayed flower senescence is also apparent in the *bos1* Crispr-Cas9 alleles. Bar = 1 cm. **(B)** Anthers of the Crispr-Cas9 knock-out alleles of *bos1* exhibited delayed dehiscence. Anthers were detached from flowers at the same developmental stage (floral stage 14). Bar = 50 μm. **(C)** The percentage of anthers that have undergone dehiscence in the indicated genotypes. Ten flowers of each genotype at the same developmental-stage were measured. The experiment was repeated twice with similar results and one representative experiment is shown. Letters above bars indicate significant differences between groups (one-way ANOVA, *P* ≤ 0.05).

### The *bos1-1* allele is a gain-of-function due to increased *BOS1* transcript levels

T-DNA insertions can result in genome structure changes or have epigenomic impacts, which may contribute to the phenotypes independent of the T-DNA insertion (Jupe et al., 2019). To comprehensively assess the genomic changes in *bos1-1*, we performed Nanopore genome re-sequencing (Brown and Clarke, 2016). This analysis identified 1173 structural variations, including larger rearrangements (>1000 bp), 13 insertions, 16 deletions, 19 duplications and 24 inversions, while no mutations were detected in the open reading frame of *BOS1* (Supplemental Table S1). Considering the work required to assess the potential role of these changes in the *bos1-1* phenotypes, we first used a strategy to test the effect of additional exon disrupting mutations in the *bos1-1* background to probe whether the *bos1-1* phenotypes were caused by increased *BOS1* transcriptional levels. With Crispr-Cas9 we introduced a second mutation in exon 2 of *BOS1* in the *bos1-1* mutant background (Fig. 3A). These intragenic double mutant alleles were assigned the designations *bos1-c4\** and *bos1-c5\** (* indicates that the allele is an intragenic double mutant in *bos1-1* background). Frame-shifts disrupting *BOS1* exon 2 were detected in these intragenic double mutants (Fig. 3B). These second mutations did not attenuate the high *BOS11* transcript levels seen in *bos1-1*, as *BOS1* transcript accumulation remained high in *bos1-c4\** and *bos1-c5\** (Fig. 3C). The characteristic *bos1-1* phenotypes were abolished in these lines: spreading cell death and *Botrytis* susceptibility in *bos1-c4\** and *bos1-c5\** were similar to wildtype (Fig. 3D and E), indicating that these exon disrupting alleles act as intragenic suppressors of *bos1-1*. Collectively, these findings demonstrate that the *bos1-1* phenotypes were caused by the alterations to *BOS1* function, rather than other genomic changes in *bos1-1*.

**Figure 3.**
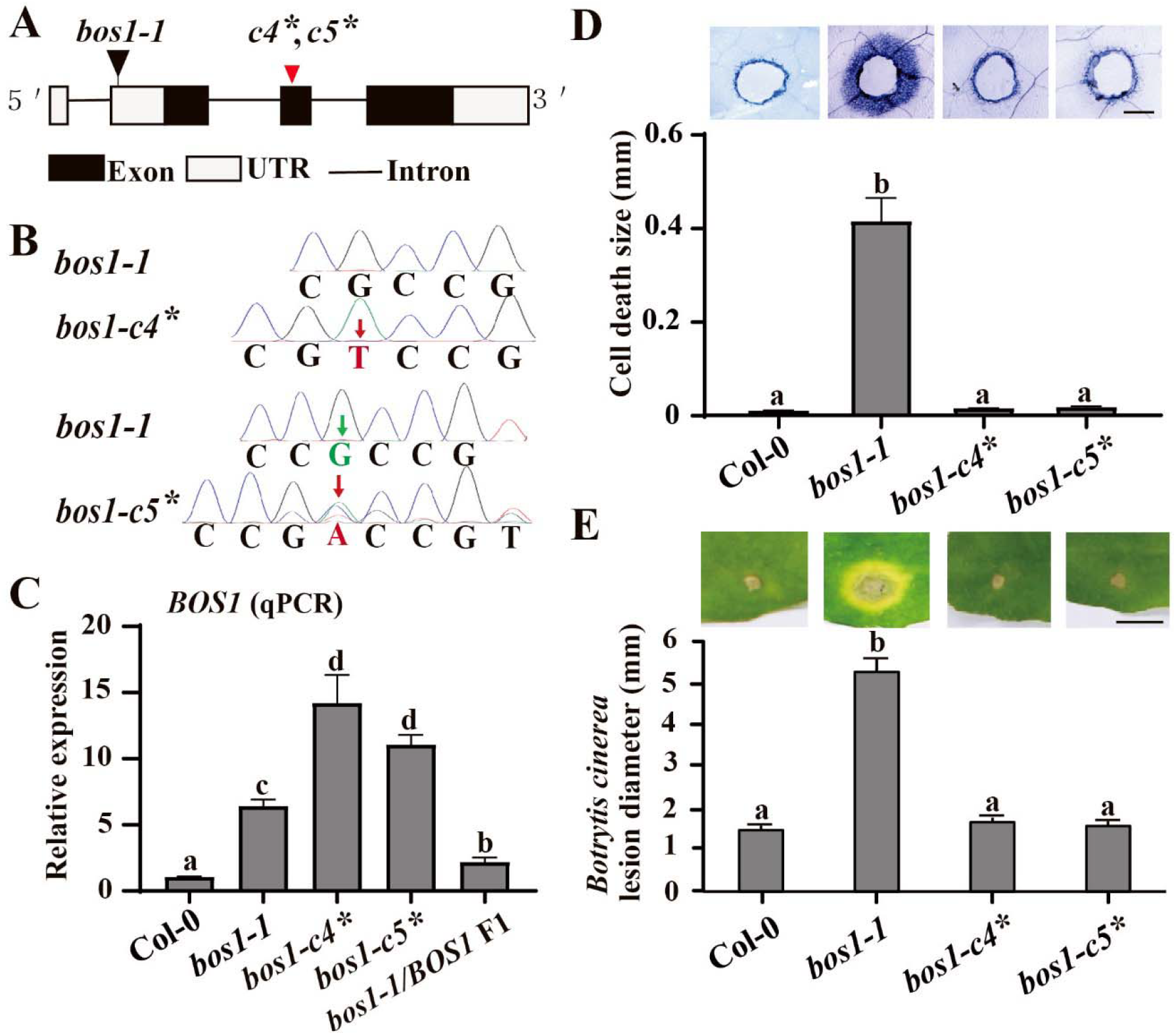
The *bos1-1* phenotypes were abolished by introduction of exon disrupting alleles in *BOS1* with Crispr-Cas9. **(A)** Schematic diagram of the new intragenic double mutants *bos1-c4\** and *-c5\** created with CRISPR-Cas9 in the *bos1-1* background, incorporating both the T-DNA insertions of *bos1-1* and frame-shifts in the second exon of *BOS1*. Black triangle indicates the T-DNA of *bos1-1*. Red triangle indicates the start of the frame-shifts of *bos1-c4\** and *bos1-c5*\* (*c4* c5\**). **(B)** Single base insertions (red characters) and deletion (green characters) were detected with Sanger sequencing. The thymidine nucleotide insertion in *bos1-c4\** is homozygous, while in *bos1-c5\** there are two changes, an insertion of an adenine nucleotide and a deletion of a guanine nucleotide. **(C)** Relative expression of *BOS1* in the indicated genotypes. Fully expanded leaves of 24-day-old plants were used for qPCR. Three biological replicates exhibited the same trends and one representative is shown. **(D)** Wounding induced cell-death spread was visualized with trypan blue staining. Representative pictures are shown to illustrate the dead tissues around the toothpick-puncture wounds. These experiments were performed three times with similar results (*n*=24 in total). Scale bar=0.5 mm. **(E)**. *Botrytis* induced lesion sizes in the indicated genotype. Statistical analysis was performed with one-way ANOVA (5 independent biological replicates; *n*=72 in total). All error bars represent the SE of means. Letters above the bars indicated significance groups (*P*<0.05, linear mixed model). Scale bar=0.5 cm.

We hypothesised that the T-DNA insertion caused altered expression of *BOS1*, which conferred the cell death phenotype in *bos1-1*. The exact site of the T-DNA insertion was unclear (Kraepiel et al., 2011). We used genome resequencing data and Sanger sequencing, which identified two adjacent T-DNAs in opposite orientations between −410 bp to −396 bp in the 5’-UTR of *BOS1* (Fig. 4A). Notably, we found a *mannopine synthase* (*MAS*) promoter in the end of each T-DNA (Fig. 4A). The *MAS* promoter is wounding inducible and controls gene expression in a bi-directional manner (Guevara-García et al., 1999). Accordingly, the expression of *BOS1* in *bos1-1* was highly wounding inducible (Fig. 4B). Each side of the *MAS* promoters resulted in expression of two detectable transcripts: the *BOS1* mRNA with a shorter 5’-UTR that was consistent with expression driven in the T-DNA and a sequence transcribed in the opposite direction derived from the T-DNA (Supplemental Fig. S2). For the *BOS1* transcript, the full coding sequence of *BOS1* was expressed and no alternative splicing events or mutations were detected (Fig. 4C; Supplemental Fig. S2). This supports that *bos1-1* phenotypes were the result of high *BOS1* transcript levels driven by the *MAS* promoter. To test this hypothesis, we transformed *pMAS::BOS1* into wildtype to test if it could confer *bos1-1* phenotypes. During generation of transgenic lines, many *pMAS::BOS1* lines exhibited pathogen susceptible phenotypes under greenhouse conditions and died after flowering (Supplemental Fig. S3). This was consistent with our previous observation that *bos1-1* did not survive under greenhouse conditions (Cui et al., 2019). In clean growth room experiments, *pMAS::BOS1* exhibited spreading cell death upon wounding, and enhanced *Botrytis* susceptibility, similar to *bos1-1* (Fig. 4D and E). Thus, the two key *bos1-1* phenoytpes were successfully reproduced by introduction of *pMAS::BOS1* to wildtype. Overall, we conclude that *bos1-1* is a gain-of-function mutant caused by *pMAS* driven expression of *BOS1*.

**Figure 4.**
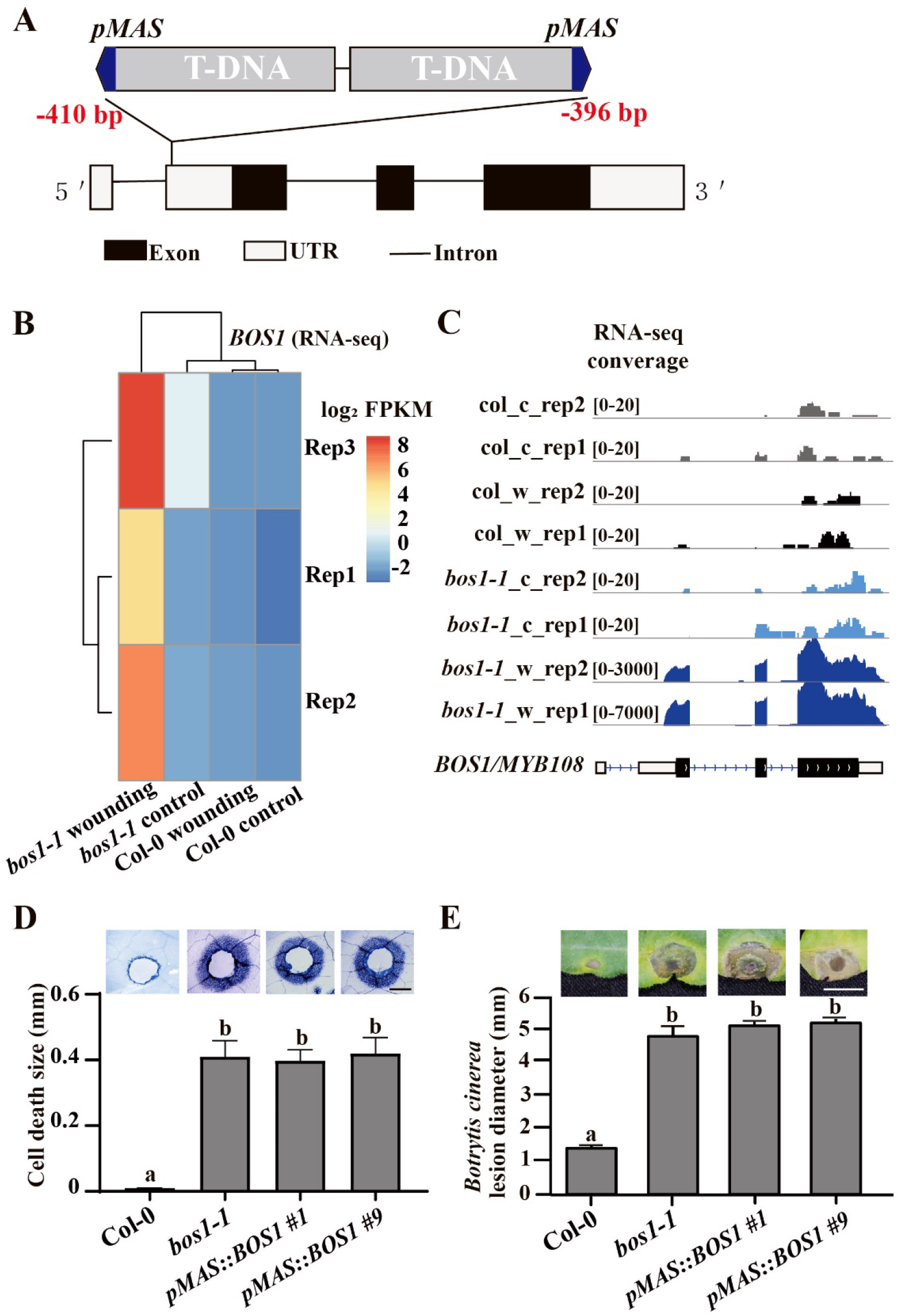
Phenotypes of *bos1-1* are caused by *MAS* promoter driven *BOS1* expression. **(A)** Schematic diagram of the T-DNA structure in *bos1-1*. Two adjacent T-DNAs were inserted in the 5’UTR of *BOS1* with *MAS* promoters indicated in blue according to re-sequencing analysis. The red numbers indicate the insertion position of the T-DNA in *bos1-1* relative to the *BOS1* start codon. **(B)** Expression profile of *BOS1* after wounding. Normalized transcript abundances of *BOS1* were calculated from RNAseq data as fragments per kilobase pair of exon model per million fragments mapped (FPKM). The log2 FPKM of indicated genotypes were used to build the heatmap. **(C)** RNA-seq reads mapped to *BOS1* genomic DNA. The entire coding sequence of *BOS1* was expressed in *bos1-1*. **(D, E)** *pMAS::BOS1* exhibited *bos1-1* mimic phenotypes upon wounding **(D)** and *Botrytis* infection **(E)** treatments. **(D)** Representative pictures and quantitative data of spreading cell death induced by toothpick-puncture wounds. Trypan blue staining was performed three times with similar results (*n*=24 in total). Scale bar=0.5 mm. **(E)** *Botrytis* induced lesion sizes are shown in the representative pictures and also as quantitative data. Statistical analysis was performed with one-way ANOVA (three independent biological replicates; *n*=24 in total). All error bars represent the SE of means. Letters above the bars indicate significance groups (*P*<0.05, linear mixed model). Scale bar=0.5 cm.

Multiple lines of evidence support a connection between *BOS1* transcript levels and *bos1-1* phenotypes: extraordinarily high *BOS1* transcript levels were detected in *bos1-1* upon wounding and *Botrytis* treatments (Fig. 3C; Mengiste et al., 2013). The extent of PCD propagation in *bos1-1* was positively correlated with the transcript levels of *BOS1*, as *BOS1/bos1-1* had lower transcript levels of *BOS1* and accordingly less cell death than *bos1-1/bos1-1* (Fig. 1C and D; Fig. 3C). To further confirm whether increased *BOS1* transcript levels enhance plant susceptibility to *Botrytis*, we constructed a series of *BOS1* overexpression lines with the *35S* promoter. We challenged these lines with *Botrytis* infection and found a positive correlation between lesion sizes and *BOS1* transcript levels (Fig. 5). This further supports that the *bos1-1 Botrytis* susceptibility was caused by increased *BOS1* transcript levels.

**Figure 5.**
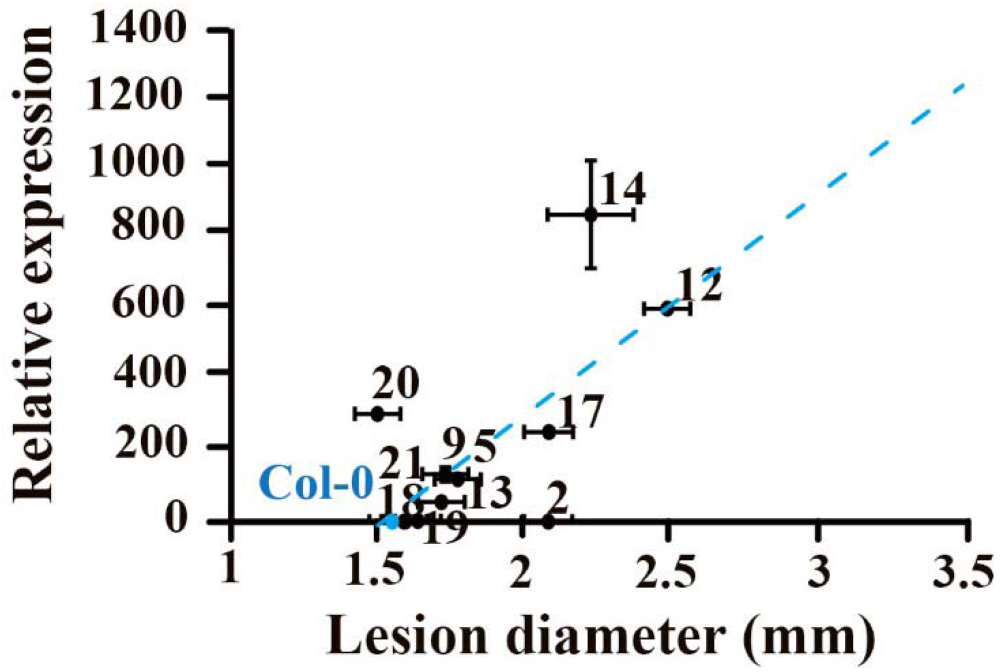
*BOS1* transcript levels in eleven *p35S::BOS1* lines were positively correlated with *Botrytis* susceptibility. The relative *BOS1* expression of the eleven independent T_2_ overexpression lines was examined by qPCR. The lesion sizes were measured as described previously. Three independent biological replicates (*n*≥48 in total) were combined and analysed. The blue dashed line indicates the correlation trend. Pearson coefficient r = 0.72, indicating a strong correlation between *BOS1* transcript levels and lesion sizes. Raw data for this figure is available in Supplemental Data Set 1.

### BOS1 in abiotic stress

*BOS1* transcript accumulation was elevated in response to multiple stresses (Supplemental Fig. S4). In order to assess the role of BOS1 in abioltic stress and hormone responses the responses to ABA, methyl viologen, and NaCl were monitored in the new loss-of-function crisper mutant alleles (Supplemental Figs. S5–S7). These experiments revealed increased sensitivity to ABA in these new mutant alleles (Supplemental Fig S5). However, mutant responses were indistinguishable from wildtype under NaCl and methyl viologen treatments (Supplemental Figs. S6–S7).

## Discussion

### Is *bos1-1* a loss-of-function or gain-of-function allele?

The inconsistent phenotypes between *bos1-1* and other *bos1* alleles were noted in previous studies; *bos1-1* exhibited no reduced fertility, in stark contrast to the clear reduction in fertility observed in T-DNA intron alleles (Mandaokar and Browse, 2009). Conversely, *bos1* T-DNA intron alleles had no pathogen phenotypes (Mandaokar and Browse, 2009; Kraepiel et al., 2011). This discrepancy only stands when *bos1-1* was interpreted as a loss-of-function mutant, which came about perhaps due to the limitation of technology in that time and a lack of other *bos1* alleles available for confirmation of phenotypes. Our data above illustrate that the cell death spread and *Botrytis* susceptibility of *bos1-1* were conferred by altered expression of *BOS1* rather than loss-of-function of *BOS1*. However, it is important to note that the original identification of *bos1-1* (Mengiste et al., 2003), used several lines of evidence, which supported well that *bos1-1* was a recessive loss-of-function mutant. This included genetic segregation and genomic complementation analyses. Importantly, the differences in procedures and conditions used in different labs may have altered the phenotypes observed. Differences in growth conditions or infection protocols can significantly influence the extent of *Botrytis* infection (Ciliberti et al., 2015; Harper et al., 1981). The key differences between our study and Mengiste et al. include fungal cultivation medium (2×V8 vs. potato dextrose broth), infection medium (Sabouraud maltose broth vs. potato dextrose broth), and the age of infected plants (3-weeks vs. 24-days). These differences may to some extent account for the different results seen between these studies. Mengiste et al. (2003) present transgenic mutant complementation data in Fig. 6C of their paper. Because of the relative lesion sizes between the complemented mutant (*bos1-1 +BOS1*) and *bos1-1* are much greater in comparison to that between complemented mutant and wild type, Mengiste et al.(2003) considered the symptoms of the complementation line as the same as wild type. However, by our own evaluation of this figure, the complemented mutant line exhibited *Botrytis* symptoms that were stronger than wild type, with larger lesion sizes and enhanced cell death around the lesion frontiers. The choice of *Botrytis* strain may also impact the extent of lesion sizes. This is well illustrated by a test of 96 diverse *Botrytis* isolates that demonstrates how different *Botrytis* strains result in contrasting symptoms (Zhang et al., 2017). The exact *Botrytis* strain was not specified in Mengiste et al., 2003. We speculate that the fungal cultivation/infection method in Megiste et al., 2003 might have led to increased contrast between the disease symptoms of *bos1-1* homozygous and *bos1/BOS1* heterozygous or the complementation lines, and obscured the intermediate phenotypes of the heterozygote or complemented line (Mengiste et al., 2003).

### Increased *BOS1* transcript levels cause *Botrytis* susceptibility and uncontained cell death

Both *pMAS* and *p35S* driven expression of *BOS1* conferred *Botrytis* susceptibility, but with some informative differences. While *pMAS* gave robust phenotypes, the *35S* promoter had outcomes that were more variable (Fig. 5). The *pMAS* promoter conferred strong wound inducible *BOS1* expression (Fig 4), which might lead to more precise *BOS1* expression at its target tissue (infection or wound sites), as compared to the general expression patterns of *p35S*. Use of *p35S* can also have unintended consequences. Multiple studies have illustrated gene silencing and integration site effects from gene overexpression using the 35S promoter (Schubert et al., 2004; Daxinger et al., 2008; Mlotshwa et al., 2010; Gelvin, 2017). To further address this, eleven *p35S::BOS1* lines were examined with *Botrytis* inoculation (Fig. 5). Most but not all of these overexpression lines exhibited enhanced susceptibility to *Botrytis*. A previous study showed that overexpression of *35S:BOS1-GUS* increased *Botrytis* resistance (Luo et al., 2010). Although the possibility that the fusion of *BOS1* to *GUS* might have altered the function of *BOS1* could not be excluded, it is not rare that some overexpression lines may exhibit phenotypes opposite to the other lines. In our study, there were also two such exceptional lines, #*2* and *#20*, among our eleven *p35S::BOS1* lines (Fig. 5). Especially, *#20* had more than 300 fold increased expression of *BOS1*, however showed slightly reduced lesion size (Fig. 5). This demonstrates the importance of evaluating many independent overexpression lines for gene function analysis.

### The *bos1* Crispr knock-outs and T-DNA alleles, but not *bos1-1*, have fertility defects

In unstressed condition, *BOS1* is mostly expressed in the cell types responsible for anther dehiscence (Mandaokar and Browse, 2009; Xu et al., 2019). Dehiscence requires properly timed PCD for pollen release (Wilson et al., 2011; Beals and Goldberg, 1997; Senatore et al., 2009). Our CRISPR knock-out lines exhibited alterations in the extent or timing of dehiscence, similar to *bos1* T-DNA intron alleles (Fig. 2B; Mandaokar and Browse, 2009; Xu et al., 2019). This suggests that *BOS1* could be required for cell death regulation in septum or stomium cells of the dehiscence zone. Thereby, the roles of *BOS1* in both stress responses and development could be unified as the requirement of *BOS1* for proper cell death regulation. The *bos1* knock-out lines have no pathogen associated phenotypes (Fig. 1C; Kraepiel et al., 2011). Only when *BOS1* transcript levels are above a certain threshold, such as in *bos1-1* after *Botrytis* infection, the cell death promoted by high expression of *BOS1* may result in altered pathogen sensitivity. *BOS1* transcript levels were elevated during multiple stresses (Supplemental Fig. S4). As this implies a role for BOS1 in abiotic defence responses, we treated *bos1-crispr* lines with ABA, NaCl and methyl viologen. The *bos1-crispr* lines exhibited enhanced sensitivity to ABA while unaltered sensitivity to methyl viologen and NaCl (Supplemental Figs. S5–S7). Further characterization of *bos1-crispr* lines to a broader range of stress and hormone treatment will help to clarify which signalling pathways are controlled by this transcription factor.

## Summary

A revaluation of previous generations of genetic tools is required (Nikonorova et al., 2018). The development of gene editing technologies allows accurate examination of gene function. These new tools facilitate re-evaluation of mutants and a refinement of our interpretation of the scientific literature (Gao et al., 2015; Westphal et al., 2008). Here, we have built upon the previous work (Mengiste et al. 2003) and demonstrated the function of BOS1 as a positive regulator of cell death. Aside from our proposed changes to some of the interpretations, the majority of this seminal paper still stands (Mengiste et al. 2003). Based on our previous publications and results here, we propose that BOS1 regulates cell death propagation signals from dying cells to neighbour cells, rather than cell death initiation. This role may be of wider interest to the plant research community and warrants further investigation.

## Materials and methods

### Cultivation conditions

Plant seedlings were transplanted to a mixture of peat and vermiculite (1:1) one week after *in vitro* growth on ½ MS medium. Plant growth conditions were 23/18 °C (day/night) temperature, 120-150 μmol m^-2^ s^-2^ light intensity, 12/12 h (light/dark) photoperiod, and 60% humidity. *Botrytis* strain Bo5.10 was cultivated on commercial medium of potato dextrose agar (PDA; P2182, Sigma-Aldrich, USA). *Botrytis* plates were kept in dark at room temperature and transferred into 4°C when conidia were produced.

### Infection and wounding assays

Fresh *Botrytis* conidia were collected with mycelium into 1/3 strength potato dextrose broth. The mixture was vortexed and filtered to remove mycelia. Conidia were suspended at a concentration of 2×10^5^ spores ml^-1^. Fully expanded leaves of 24-day old plants were inoculated with 3 μl conidia solution. Plants were covered with a transparent plastic lid to keep 100% humidity. Symptoms were photographed at 3 days post inoculation (dpi). Wounding was conducted with a toothpick by puncturing fully expanded leaves of 23-day-old plants. Wounding-induced cell death was visualized by trypan blue staining with wounded leaves collected at 4 days post wounding (dpw). Both lesion sizes and wounding-induced cell death were measured by using ImageJ (http://rsb.info.nih.gov/ij/). The basic experimental procedures of cell death staining were the same as in our previous publications (Cui et al., 2013, 2019).

### Seedling growth assays

For the ABA and NaCl treatments, sterilized seeds were sown on ½ MS media containing ABA or NaCl with indicated concentrations. The root lengths were photographed at nine days after sowing, and measured by using ImageJ (http://rsb.info.nih.gov/ij/). For methyl viologen treatment, seeds were germinated on control plates and four-day-old seedlings were transferred to media with indicated concentrations of methyl viologen. Photos were taken at 15 days after transplanting.

### Cloning procedures

The genomic DNA of *BOS1* was cloned into the vector pGWB412 to construct the *p35S::BOS1* plasmid. The *MAS* promoter was cloned with template DNA from the *bos1-1* mutant, and then replaced the 35S promoter of *35S::BOS1* to create *pMAS::BOS1*. For Crispr-Cas9 knock-out alleles, guide RNA (gRNA) targeting the first and second exons of *BOS1* were integrated into pCBC_DT1DT2 and then into the final vector pHEC401 according to (Xing et al., 2014). Vectors were transformed into the indicated plants *via* Agrobacterium strain GV3101. The primers used in this study are listed in Supplemental Table2.

### Transformation procedures

The *bos1-1* mutant is not amenable to transformation with Agrobacterium. The Agrobacterium transformation of *bos1-1* was performed in labs in Helsinki, Finland and Hangzhou, China. All *bos1-1* plants died before seed set because of the spreading cell death trigged from Agrobacterium infection. To overcome this limitation, our strategy was that the Crispr-Cas9 vectors were first transformed to wildtype, and then introduced to *bos1-1* via crossing of *bos1-1* and the transformed wildtype.

### Genome re-sequencing

Genomic DNA of *bos1-1* was extracted and sequenced by the Biomarker Technologies Corporation (Beijing, China) following the standard procedures of Oxford Nanopore Technology sequencing (Deamer et al., 2016). Sequence depth was 129x, 99.77% of 24.37 Gb clean data mapped properly to the *Arabidopsis* genome (TAIR10). The raw data has been deposited to NCBI (PRJNA728243). Structural variations were analyzed with Sniffles (Sedlazeck et al., 2018).

### RNA-sequencing

Fully expanded leaves of 23-day-old plants were punctured with bunched toothpicks, and collected after three days. Unwounded plants were used as control. RNA was extracted with TRIzol reagent (Invitrogen, Carlsbad, CA, USA), library construction and sequencing were carried out in LC-BIO Bio-tech Ltd with Illumina Hiseq 4000. Raw reads were filtered and aligned to the *Arabidopsis* genome (TAIR10) using the hisat2 (v2.1.0) (Kim et al., 2015). To identify the transcripts adjacent to *pMAS*, we first obtained the conjoined sequence of the T-DNA and *BOS1* genome sequence from the *bos1-1* mutant resequencing analysis, verified the sequence with Sanger sequencing, and then used the combined sequence as reference for read mapping. The RNA seq raw data has been deposited to NCBI (PRJNA728243). Normalized transcript abundances of *BOS1* were calculated as fragments per kilobase pair of exon model per million fragments mapped (FPKM) with Cufflinks (Trapnell et al., 2010). For real-time quantitative PCR (qPCR), leaves of 23-day-old plants were used for RNA extraction and reverse transcription. The raw cycle threshold values were analyzed with Qbase+ (Biogazelle; Hellemans et al., 2007) with the reference genes *Actin2, PP2AA3*, and *Actin8*.

### Statistical analysis

The experiments of *Botrytis* inoculation and wounding treatments were performed at least three times. Lesion sizes and cell death spread were analyzed with scripts in R (version 3.0.3). Briefly, combined experiments were subjected to a linear mixed model with the nlme package with fixed effect for genotypes, treatments and a random effect for biological repeats. Multcomp package were used to estimate the contrasts and single-step *P*-value correction were used to estimate the *P*-value. Pearson coefficient calculations in R were used to support the strength of correlation.

### Accession Numbers

Gene identifiers for *Arabidopsis* BOS1/MYB108 (AT3G06490), Actin2 (AT3G18780), PP2AA3 (AT1G13320), Actin8 (AT1G49240). New sequencing data, including *bos1-1* resequencing data and RNA-seq data can be found at the NCBI SRA (PRJNA728243).

## Supporting information

Identification of genomic changes in bos1-1, identified through genome re-sequencing.

Primers used in this study

Raw data for Figure 5, including lesion sizes and BOS1 transcript levels.

## Supplemental Data

The following materials are available in the online version of this article.

**Supplemental Table S1**: Identification of genomic changes in *bos1-1*, identified through genome re-sequencing.

**Supplemental Table S2**: Primers used in this study.

**Supplemental Data Set 1**: Raw data for Figure 5, including lesion sizes and *BOS1* transcript levels.

## Acknowledgments

This work supported by the Natural Science Foundation of Zhejiang Province (grant no. LY22C160005); the National Natural Science Foundation of China (grant no. 31700224 and 31871233); No conflict of interest declared.

## Supplemental Figures

**Supplemental Figure S1.**
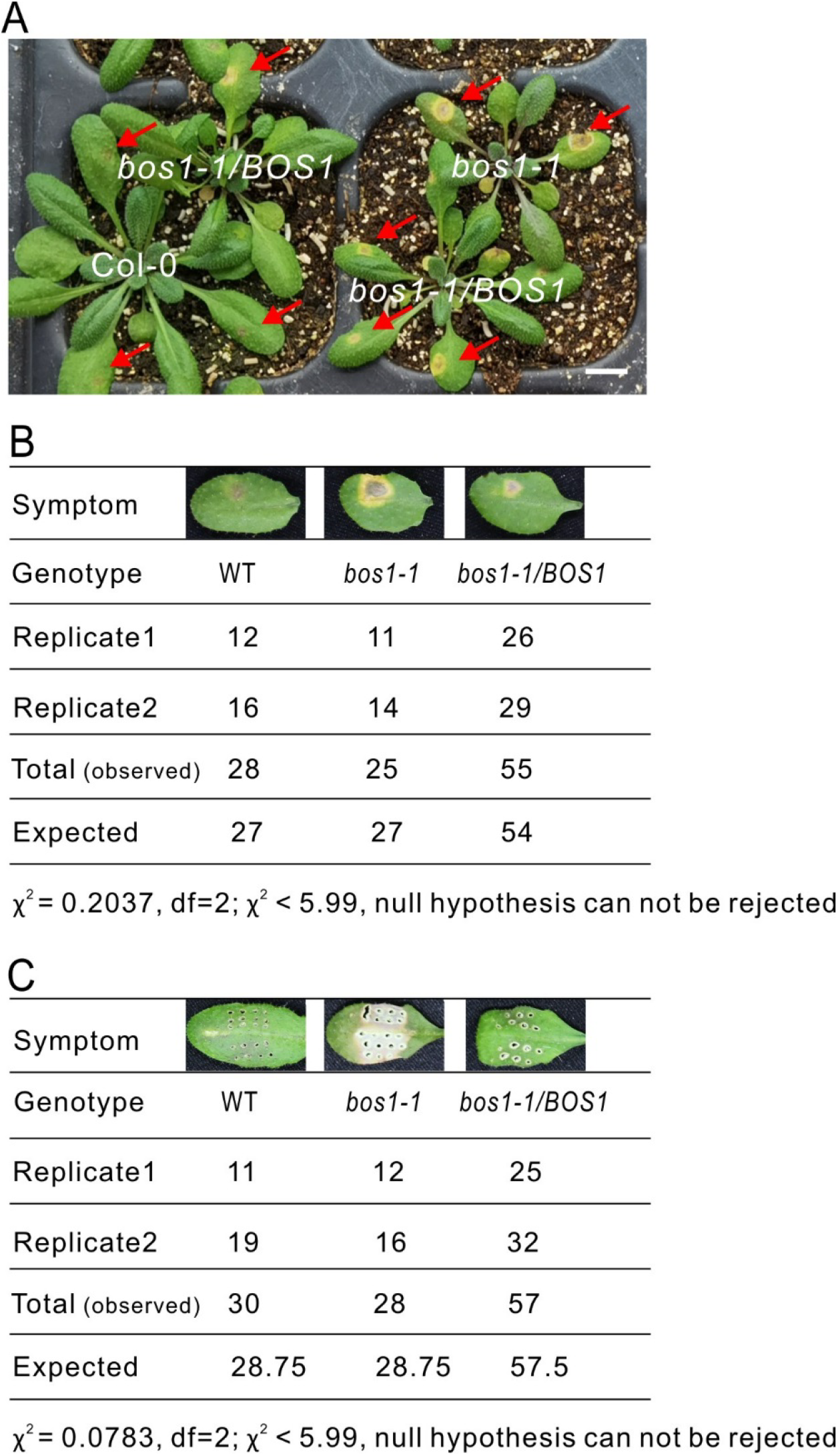
Phenotype segregation in F_2_ plants in response to *Botrytis* infection and wounding treatments. (A) Plant symptoms induced by *Botrytis* inoculation. Representative plants of known genotypes, which were confirmed by PCR. Bar = 1 cm. (B-C) The number of F_2_ *bos1-1/Col-0* individuals exhibiting the indicated symptoms upon *Botrytis-* (B) and wounding- (C) treatments. The genotypes of several plants were confirmed by PCR and are presented as representative symptoms. F_2_ individuals with similar symptoms were counted and the number of individuals in each category are listed. A model with a co-dominant effect of *bos1-1* was used as the null hypothesis for the χ^2^-test.

**Supplemental Figure S2.**
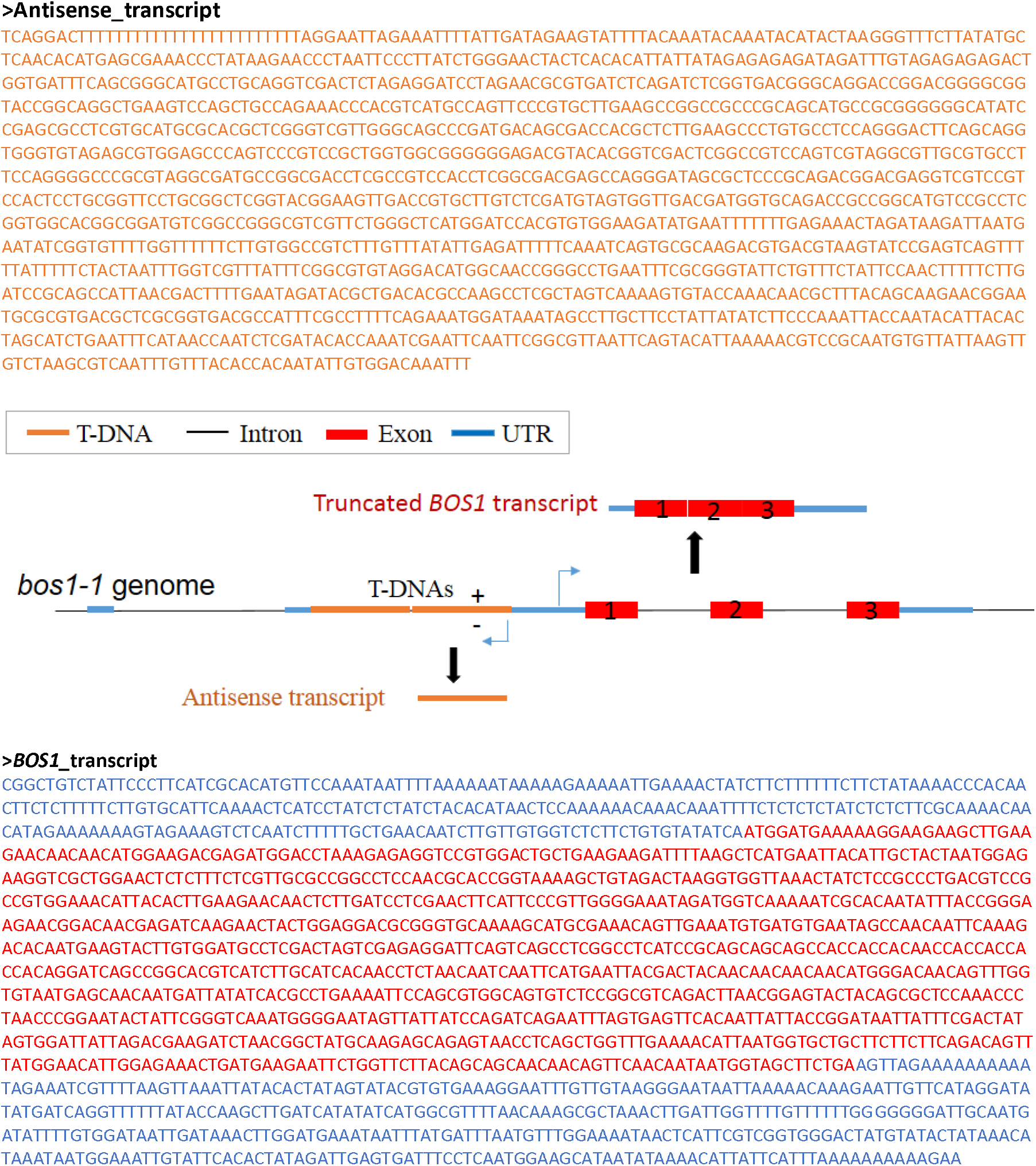
Illustration of the transcripts at the *BOS1* locus in *bos1-1* mutant. Two transcripts were found in the RNA-seq analysis. One is the truncated *BOS1* transcript with a shorter 5’-UTR, and the other is an antisense transcript from the T-DNA.

**Supplemental Figure S3.**
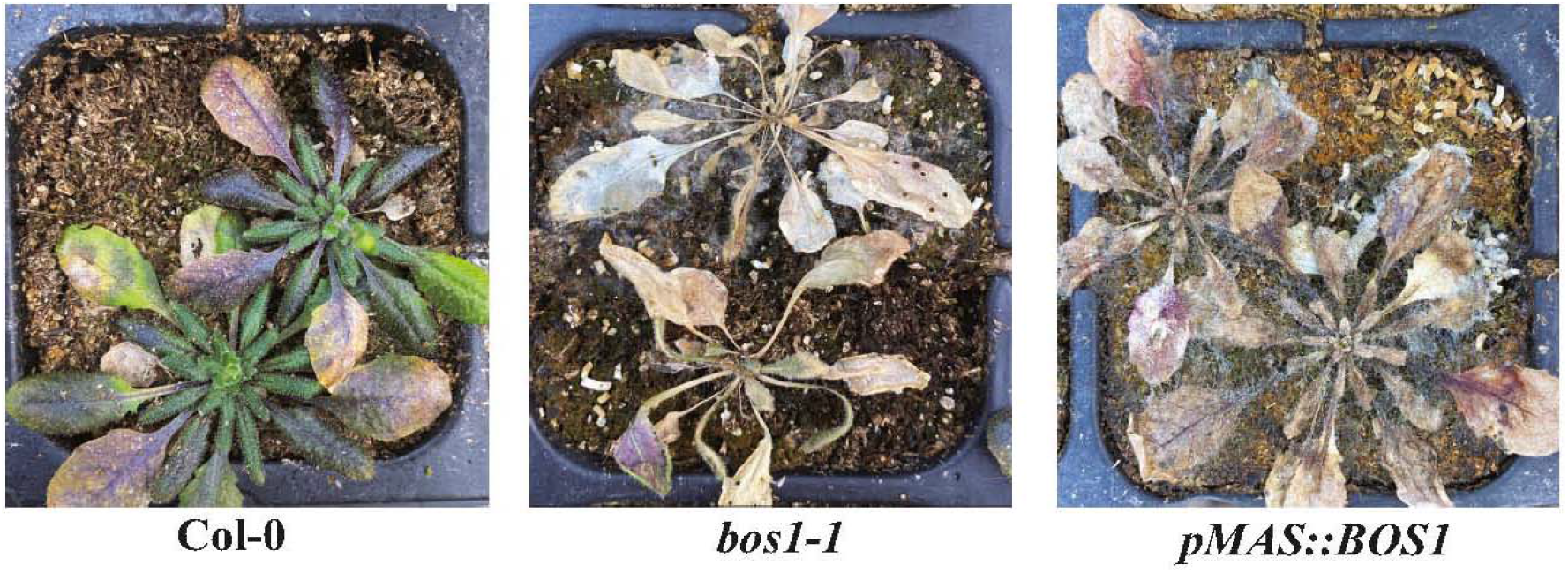
The *pMAS: BOS1* lines were more sensitive to pathogens under standard greenhouse conditions. Plants grown in the greenhouse without fungicide application. Many *pMAS: BOS1* lines were infected and died before setting seed. Representative plants of the indicated genotypes are shown.

**Supplemental Figure S4.**
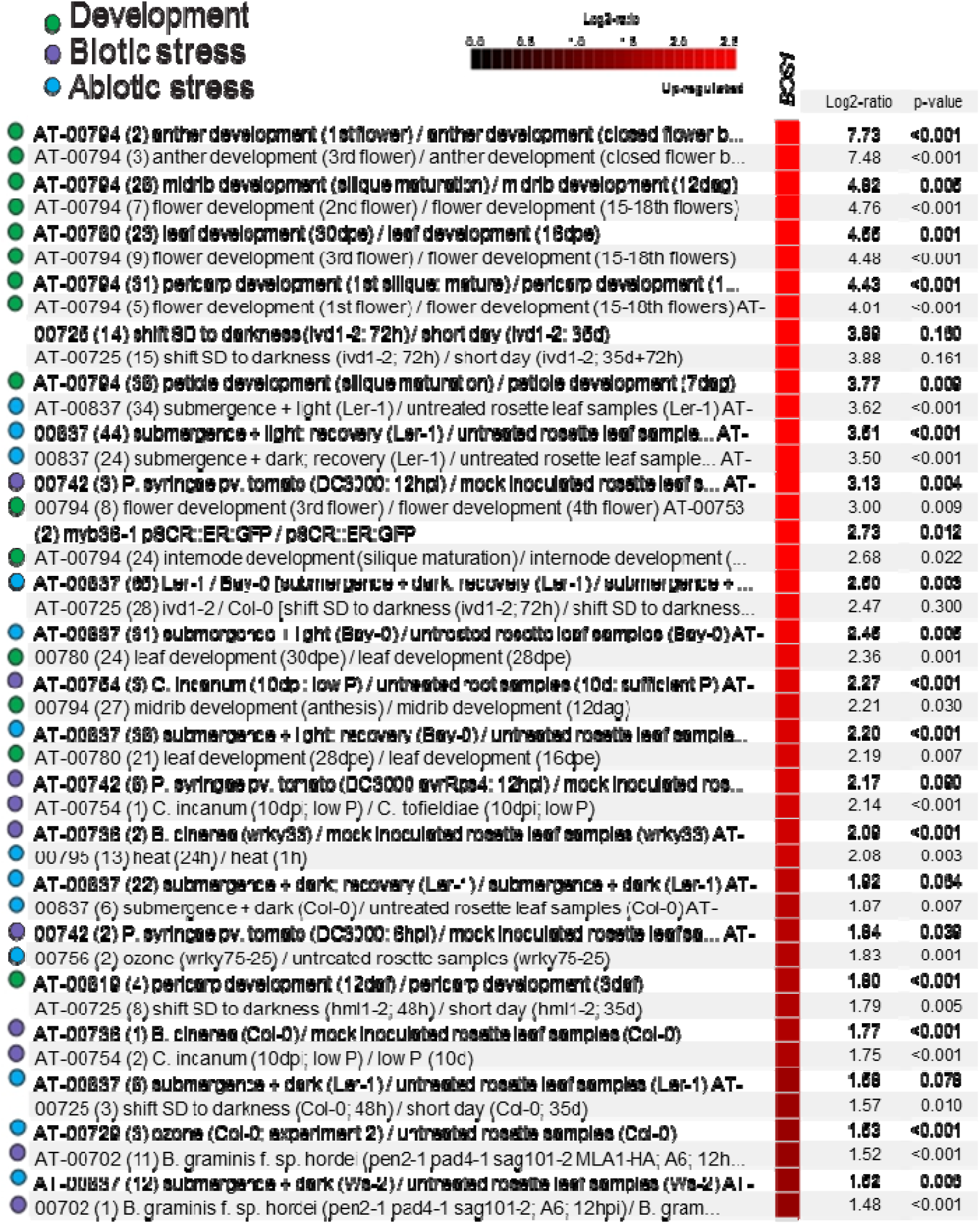
*BOS1* transcript levels in publicly available Arabidopsis RNAseq data. The Genevestigator perturbation tool was used to identify experiments with highest up-regulation of BOS1 transcript levels (Hruz et al., 2008). The identification number for each experiment refers to the identifier in the Genevestigator database.

**Supplemental Figure S5.**
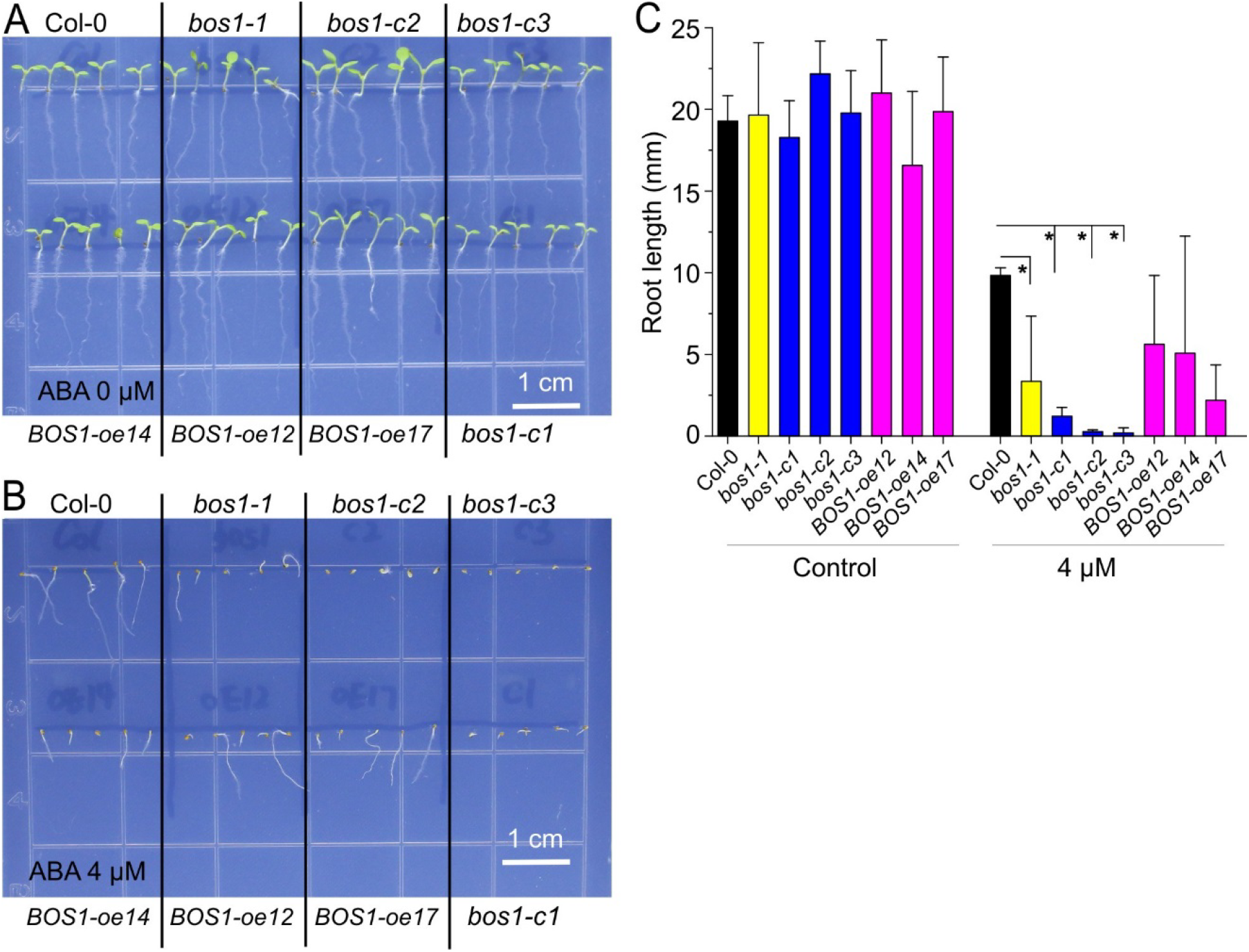
The *bos1-crispr* loss-of-function mutants exhibited enhanced ABA sensitivity. (A-B) Symptoms of the plants in response to ABA at the indicated concentrations. These experiments were repeated twice with similar results. Bar = 1 cm. (C) Pooled quantitative data of the root lengths of two independent biological repeats. Stars indicated the significant different groups (p < 0.05; *t*-test).

**Supplemental Figure S6.**
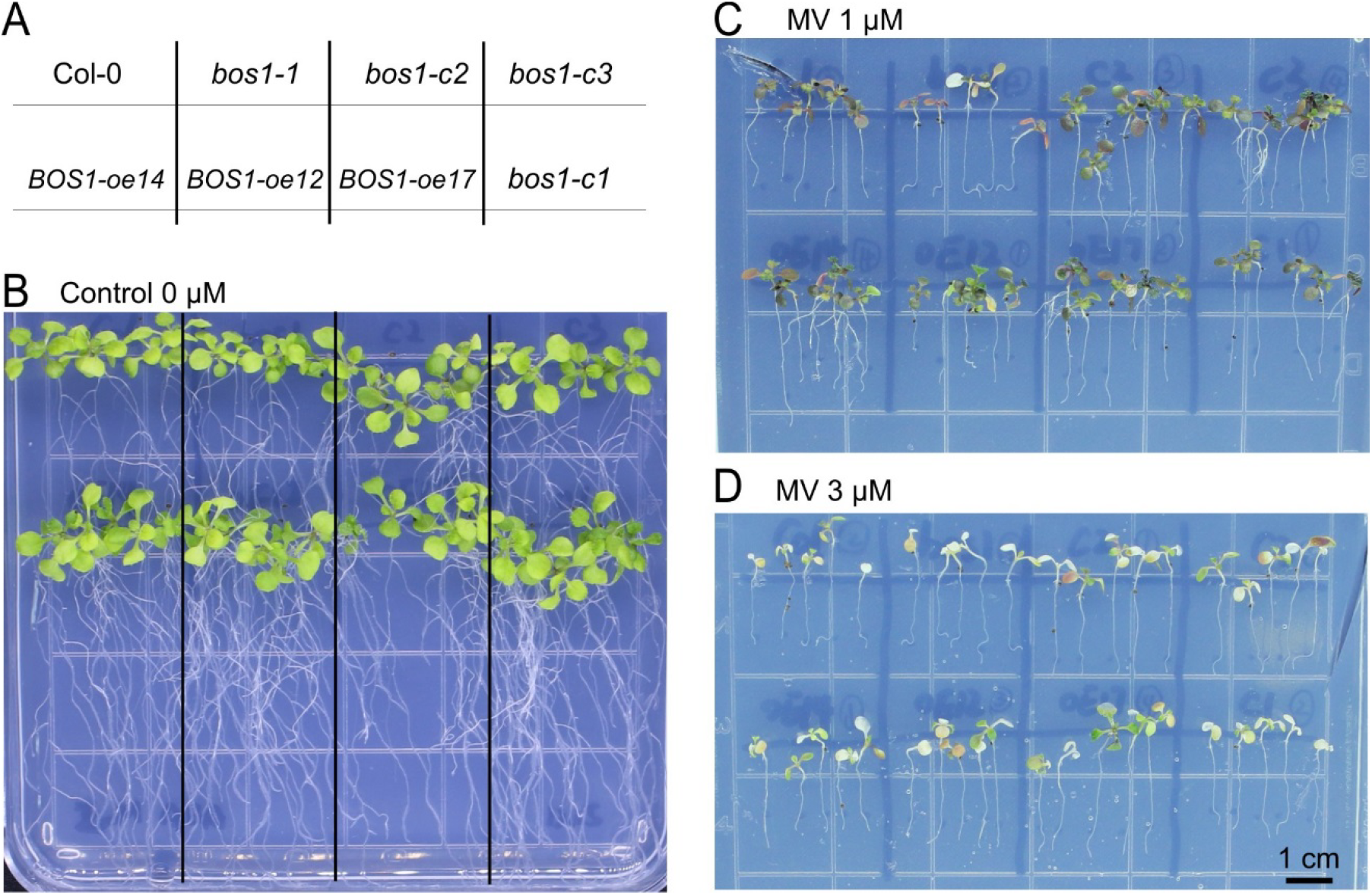
The *bos1*-crispr loss-of-function mutants exhibited unaltered sensitivity to methyl viologen. (A) Illustration of the plant genotypes (B-D) Symptoms of the plants in response to methyl viologen at the indicated concentrations. These experiments were repeated twice with similar results and one representative experiment is shown. Bar = 1 cm.

**Supplemental Figure S7.**
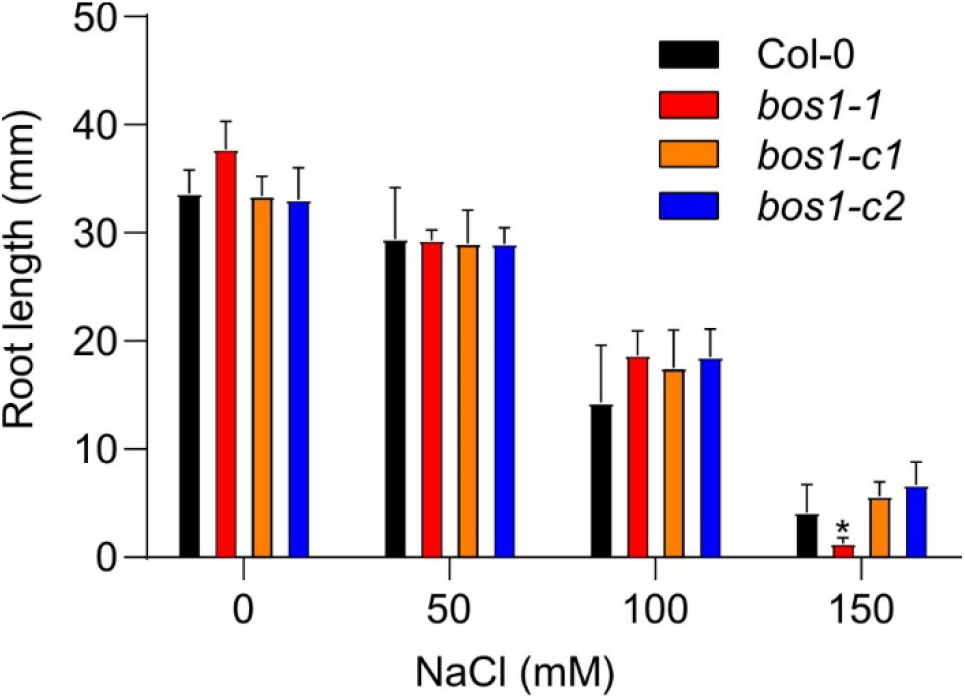
The *bos1*-crispr loss-of-function mutants exhibited unaltered NaCl sensitivity. The root length of the indicated genotypes were measured on the 9^th^ day. These experiments were repeated twice with similar results (n=20 in total). Stars indicate the groups that are significantly different (p < 0.05; *t*-test).

