## Supplementary material for "BOS1 is a positive regulator of wounding induced cell death and plant susceptibility to *Botrytis*": Primers used in this study

Table S2 Primers used in this study.

| Primer | Sequence |
| --- | --- |
| pGWB412-*BOS1*-F | GGGGACAAGTTTGTACAAAAAAGCAGGCTTC  ATGGATGAAAAAGGAAGAAGCTTG |
| pGWB412-*BOS1*-R | GGGGACCACTTTGTACAAGAAAGCTGGGTTTTT  CAGAAGCTACCATTATTGTTGAAC |
| pGWB412-*MAS*-F | TTGTAAAACGACGGCCAGTGCCAAGCTTTT  CAGCAGGTGGGTGTAGAGC |
| pGWB412-*MAS*-R | GATGCAAATTGTCCTAAAGGGTC |
| *BOS1T*-F | GTCTTAACCTTCACGCACATAAAA |
| *BOS1T*-R | CGCAACGAGAAAGAGAGTTCC |
| *BOS1*-TDNA-F | GCGTAGTTGCTTTGAGCGTGG |
| *BOS1*-TDNA-R | GGCAAACGCTTTACGCTGAAAC |
| *BOS1*-E_2_E_3_-F | GGGAAGAGCACTAACTCAATGG |
| *BOS1*-E_2_E_3_-R | CTGACTGAATCCTCTCGACTAG |
| *BOS1-*P_3_E_1_-F | ATTACGGCTGTCTATTCCCTTC |
| *BOS1-*P_3_E_1_-R | CATACCGGCGCAACGAGAAA |
| *BOS1*-qF-1 | ACGGCTATGCAAGAGCAGAGTAAC |
| *BOS1*-qR-1 | ACTGTCTGAAGAAGAAGCAGCAC |
| *AtACTIN2*-qF | GGTAACATTGTGCTCAGTGGTGG |
| *AtACTIN2*-qR | AACGACCTTAATCTTCATGCTGC |
| *AtACTIN8*-qF | ATGACTCAGATCATGTTTGAGACC |
| *AtACTIN8*-qR | TCAGTAAGGTCACGACCAGCAA |
| *BOS1*-Ex1-F0 | TGTACTAATGGAGAAGGTCGCGTTTT  AGAGCTAGAAATAGC |
| *BOS1*-Ex1-BsF | ATATATGGTCTCGATTGTACTAAT  GGAGAAGGTCGCGTT |
| *BOS1*-Ex2-F0 | TGTCCGCCCTGACGTCCGCCGGTTTT  AGAGCTAGAAATAGC |
| *BOS1*-Ex2-BsF | ATATATGGTCTCGATTGTCCGCCC  TGACGTCCGCCGGTT |
| *BOS1-*Ex3*-*R0 | AACGCGTCCTCCAGTAGTTCTTCAA  TCTCTTAGTCGACTCTAC |
| *BOS1-*Ex3*-*RsR | ATTATTGGTCTCGAAACGCGTCC  TCCAGTAGTTCTTC |
| *BOS1*-Pr3*-*R0 | AACCAACAAGATTGTTCAGCAAC  AATCTCTTAGTCGACTCTAC |
| *BOS1*-Pr3*-*BsR | ATTATTGGTCTCGAAACCAACA  AGATTGTTCAGCAAC |
